# Development of novel temporal beta-diversity indices for assessing community compositional shifts accounting for changes in the properties of individuals

**DOI:** 10.1101/2022.04.13.488244

**Authors:** Ryosuke Nakadai

## Abstract

Revealing the patterns of temporal biodiversity changes and deciphering the connection between individual life histories and large-scale ecological patterns is essential for improving the mechanistic understanding of macroecology. However, this is challenging because the relationship between individual life history and biodiversity remains unclear. In the present study, in order to link the individual life history and community-level phenomena, I developed novel indices that allow the evaluation of community compositional shifts over time by explicitly considering the contributions of the life histories of individuals (i.e. growth, mortality, and recruitment) in a community. These novel indices are quantitative extensions of the individual-based temporal beta-diversity indices which can include information on individual sizes. The indices were applied to a subset of data from the US Forest Inventory and Analysis database for the state of Rhode Island, USA, to identify changes in the contribution of individual life histories to biodiversity change. The results of this study represent methodological progress in community ecology and macroecology, as well as a conceptual advancement in bridging studies on biodiversity with those on individual life history and physiology. The individual-based diversity indices developed here pave the way for individual-based biodiversity science, which may facilitate the understanding of the effects of climate change across different hierarchies of biological organisation.

## 1. Introduction

An individual cannot exist in two places simultaneously, but it can persist in the same habitat over time (Magurran et al., 2019). This feature of individuals is a major difference between time and space in community ecology. In a temporal context, individual turnover between different species is directly linked to compositional shifts in a community (Magurran et al., 2019, Nakadai 2020). Therefore, individual persistence and turnover are important factors for measuring temporal compositional shifts in biodiversity, termed temporal beta diversity (Gotelli et al., 2017; Magurran, 2011, Tatsumi et al., 2021). For example, individuals of long-lived species (e.g. trees and vertebrates) can persist in the long term, whereas individuals of short-lived species are replaced by conspecifics or individuals of other species (Nakadai, 2020, 2021). Nakadai (2020) proposed the concept of ‘individual-based beta diversity’ to bridge the gap between biodiversity and individual information and provided novel indices to evaluate individual turnover and compositional shifts based on comparisons between two periods at a given site. Specifically, Nakadai (2020) developed an individual-based temporal beta-diversity approach for the Bray– Curtis dissimilarity index (i.e. an abundance-based temporal beta-diversity index) and accounted for the contribution of both individual persistence and turnover to the compositional shifts in biodiversity over time (Nakadai, 2020, 2021).

Climate change is known to directly affect not only biodiversity, but also the individual life histories of organisms (Brienen et al., 2020; Searle & Chen, 2018). Recent studies have shown that climate warming will accelerate the growth and shorten the life spans—which are determined by physiological processes—of individual trees (Körner, 2017; Munné-Bosch, 2018; Searle & Chen, 2018) and increase the amount of carbon stocks in terrestrial ecosystems (Brienen et al., 2015). Such changes in individual life histories potentially affect the compositional shifts of local communities and, consequently, global biodiversity. While understanding the connection between individual physiology and large-scale ecological patterns is essential for a mechanistic understanding of macroecology (Brown, 1995, 1999; Brown et al. 2004; e.g. metabolic theory), this is challenging because the connection between individual life history and biodiversity remains unknown. Nevertheless, to comprehensively understand the effects of climate change, it is essential to simultaneously capture changes in individual properties, individual turnover, and compositional shifts in biodiversity.

Despite these requirements, previous approaches to individual-based temporal beta diversity (Nakadai, 2020, 2021) have only considered the presence or absence of each individual (e.g. qualitative properties such as ‘dead’ or ‘alive’), and the effects of property changes in each individual (e.g. individual growth) on biodiversity over time have been neglected. Therefore, the aim of this study was to develop new individual-based temporal beta-diversity indices that can assess compositional shifts over time by explicitly including property changes in each individual in a community. Hereafter, I refer to the indices developed by Nakadai (2020) and the novel indices introduced in the present study individual-based ‘qualitative’ (‘QL’) indices and individual-based ‘quantitative’ (‘QT’) indices, respectively. By introducing quantitative individual information (e.g. size and weight), these novel indices can evaluate the contribution of individual growth and the differences between large and small individuals. After briefly reviewing the history of pairwise beta-diversity indices, I extended individual-based QL temporal beta-diversity indices to QT indices. I then applied these indices to subset data from the US Forest Inventory and Analysis (FIA) database for the state of Rhode Island in the USA (Stanke et al., 2020, Burrill et al., 2021) as a case study to demonstrate their application.

## 2. Materials and methods

### 2.1. Conventional incidence- and abundance-based indices and recently developed individual-based qualitative (QL) temporal beta-diversity indices

Incidence-based and abundance-based beta-diversity indices have been previously proposed (Baselga, 2013). For example, the widely used Bray–Curtis dissimilarity index (Odum, 1950) is an abundance-based extension of the Sørensen index (Baselga, 2013; Legendre & Legendre, 2012; Sørensen, 1948). Incidence-based methods target the number of species, whereas abundance-based methods mainly target the number of individuals and, sometimes, area, biomass, and weight (e.g. Li et al., 2016). Conventional beta-diversity indices are based on intersection (*A, a*) and relative complement (*B, b, C, c*) components. For context, the Sørensen dissimilarity index (*d*_*sor*_; Sørensen, 1948) is formulated as follows:

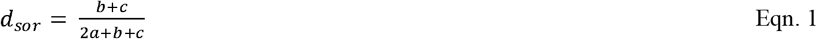

where *a* is the number of species common to both temporal units; *b* is the number of species that occur in the first but not the second temporal unit; and *c* is the number of species that occur in the second but not the first temporal unit (Legendre, 2019). Analogously, the Bray–Curtis dissimilarity index (*d*_*BC*_; Odum, 1950) is formulated as follows:

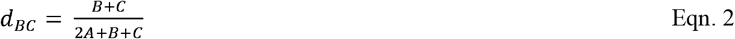

As a premise for explanation, assume that there are two sets of community data along the time axis—communities at time *j* and time *k* (abbreviated as T*j* and T*k*: *j* < *k*), respectively. The abundance-based components are as follows:

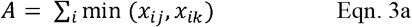

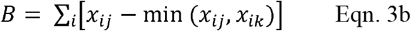

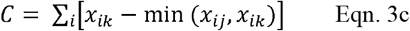

where *x*_*ij*_ is the abundance of species *i* at temporal unit *j*, and *x*_*ik*_ is the abundance of species *i* at temporal unit *k*. Therefore, *A* is the total number of individuals of each species shared by units *j* and *k*, and *B* and *C* are the total numbers of individuals of each species unique to units *j* and *k*, respectively (Odum, 1950, Legendre, 2019).

Nakadai (2020) introduced the categorisation of the community dynamics component into persistence (*p*), mortality (*m*), and recruitment (*r*) in individual-based temporal beta diversity. *m* refers to the number of individuals occurring at T*j* that die before T*k*, representing the total number of individuals that died during the time between T*j* and T*k. r* identifies the number of individuals that were not yet alive at T*j* (or were not yet in the size class counted, e.g. diameter at breast height (DBH) ≥ 1 cm), but that were alive and counted at T*k*, representing the total number of individuals recruited between T*j* and T*k*. Component *p* refers to the persistence of individuals from T*j* to T*k*. Note that, unlike previous studies (Nakadai, 2020, 2021; Fig. 1a), I use lower-case letters for the components of individual-based qualitative indices (i.e. *p, m*, and *r*) to distinguish the components of the individual quantitative indices described later. Thus, the abundances are calculated as follows:

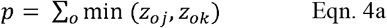

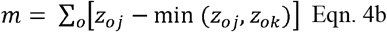

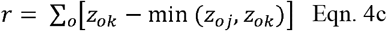

where *z*_*oj*_ is individual *o* at time T*j*, and *z*_*ok*_ is individual *o* at time T*k*. Both *z*_*oj*_ and *z*_*ok*_ have values of one or zero (presence and absence, respectively). Therefore, *p* is the total number of individuals present at both T*j* and T*k*, and *m* and *r* are the numbers of individuals unique to T*j* and T*k*, respectively (Nakadai, 2020, 2021; Fig. 1a).

**Fig. 1.**
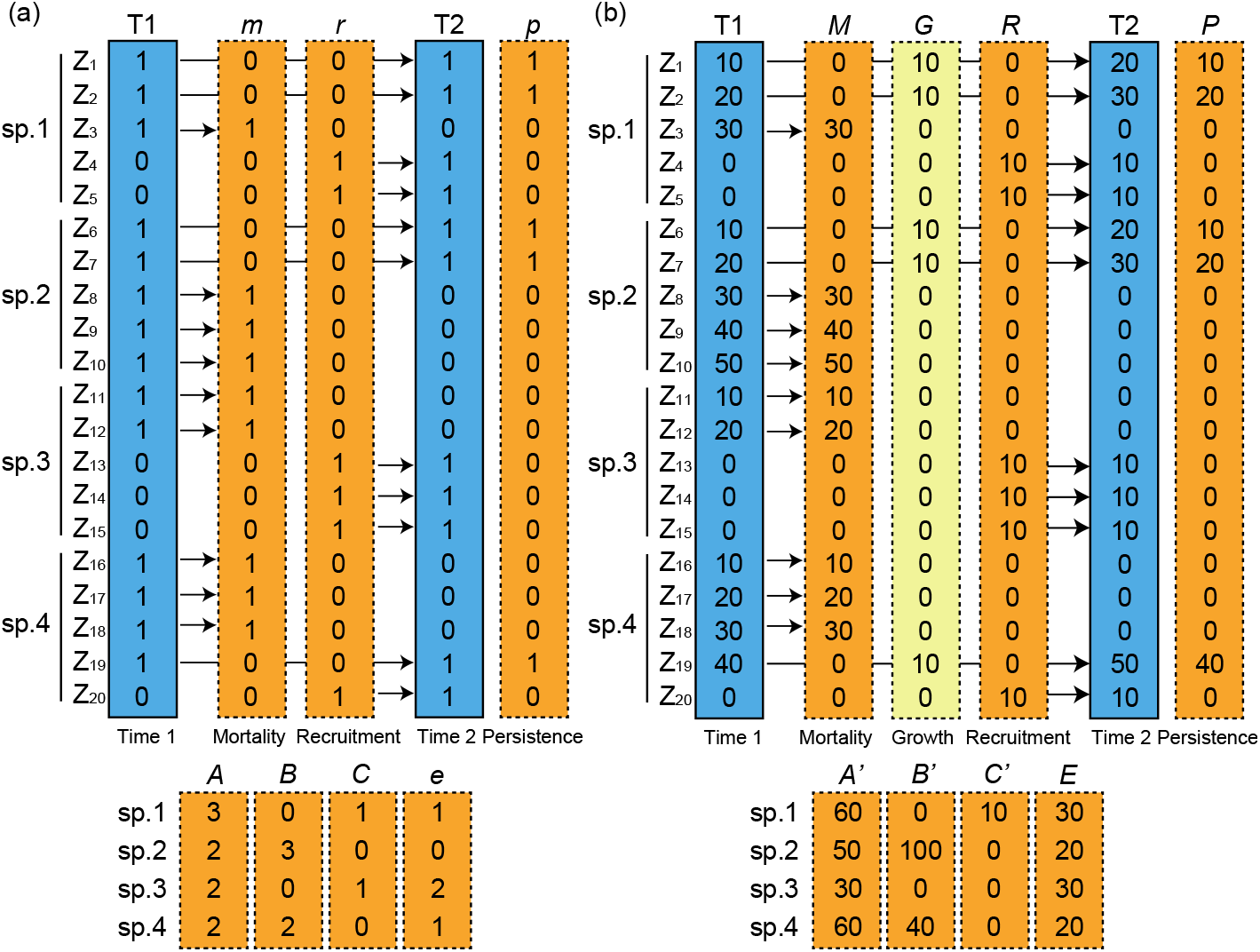
Schematic representation of individual-based temporal beta diversity for both qualitative (a) and quantitative (b) components. T1 and T2 refer to the communities at times 1 and 2, respectively. Both communities T1 and T2 comprise four species (sp. 1, 2, 3, and 4), and all individuals are identified separately (*Z*_*o*_) in both cases. The arrows indicate the temporal trajectory for each individual. In the qualitative components (a), *m* refers to the number of individuals in existence at T1 but dead before T2, thereby separately identifying the deaths of individuals during the period between T1 and T2; *r* refers to the number of individuals that were not yet alive at T1 but were present at T2, thereby identifying the recruitment of individuals during the period between T1 and T2; *p* refers to the number of individuals that persisted from T1 to T2, thus both T1 and T2 = 1; *e* refers to the number of individuals that did not contribute to the apparent compositional shift among mortality or recruitment of individuals, which is calculated as *m*–*B* or *r*–*C*. In the qualitative components (b), *M* refers to the total measures of individuals occurring at T1 that died before T2, thereby representing the total measures of the mortal individuals during the time between T1 and T2; *R* refers to the total measures of individuals that were not yet alive at T1 but were alive and counted at T2, thereby representing the individuals recruited between T1 and T2. Persistent individuals are qualitatively identical, having alive (or present) status between T1 and T2, but rarely maintain the same status with respect to size and weight (e.g., no size change). Therefore, changes in individual sizes occur between T1 and T2. The quantitative indices consider the contribution of changes in individuals’ sizes, mainly targeting individual growth. Specifically, for persistent individuals, the original and altered total measures are partitioned into persistent (*P*) and growth (*G*) components, respectively. *E* refers to the total basal area that did not contribute to the apparent compositional shift among mortality, recruitment of individuals, or growth of individuals, which is calculated as *M+G*_*de*_–*B’* or *R +G*_*in*_ –*C’*.

The individual-based temporal beta-diversity index for Bray–Curtis dissimilarity (*d*_*mr*_) can be formulated (Nakadai 2020) as follows:

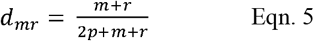

In some cases, there is a turnover of different individuals belonging to the same species, even when the community composition is stable over time. Therefore, the components that change over time (*m* and *r*) are reorganised into two further components, namely compositional equilibrium (*e*) and shift (*B* and *C*) (Nakadai, 2020, 2021). Here, I define compositional equilibrium as a turnover component without apparent compositional shift in two time steps, thereby contributing to the dynamic equilibrium. Component *m* can be partitioned into two additive components—the lost individuals that contribute to compositional shifts in a community (*B*) and the lost individuals replaced by conspecifics that contribute to an equilibrium in community composition (*e*_*loss*_) (Fig. 1a). Similarly, component *r* can be partitioned into two additive components—gained individuals that contribute to compositional shifts in a community (*C*) and gained individuals that replace conspecifics, thereby contributing to an equilibrium in community composition (*e*_*gain*_) (Fig. 1a). By definition, the numbers of *e*_*loss*_ and *e*_*gain*_ individuals are identical; thus, I replaced both *e*_*loss*_ and *e*_*gain*_ with the coefficient *e* (Nakadai, 2020, 2021). The persistence component (*p*) is the shared (apparent non-changed) component (*A*) minus the equilibrium (changed, but apparent non-changed) component (*e*) of community composition. The above descriptions are summarised below:

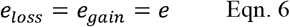

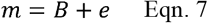

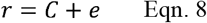

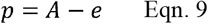

Furthermore, components *m* and *r* can be analysed in the same manner as the community data (Nakadai, 2020). Therefore, the Bray–Curtis dissimilarity between communities can be calculated based on *m* and *r* (Fig. S1d), as follows:

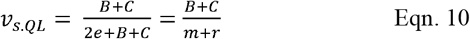

where *v*_*s.QL*_ indicates the speed of compositional shifts in a community relative to the total speed of individual turnover associated with *m* and *r*. In other words, *v*_*s.QL*_ indicates total changes in community composition divided by total changes in the number of individuals. It is possible to calculate *v*_*s.QL*_ as *d*_*BC*_ divided by *d*_*mr*_ as:

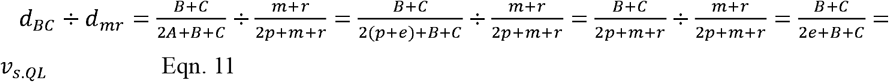

### 2.2. Novel individual-based “quantitative (QT)” beta-diversity indices

I developed individual-based QT components and novel indices to analyse community datasets over time using individual property information (e.g. size and weight). The relationship between the QL and QT indices is similar to that between the incident-based and abundance-based indices. In other words, individual-based QT components are calculated as individual-based QL weighted by individual sizes. As previously noted, the abundance targeted by abundance-based indicators is not necessarily the number of individuals but can also include area and volume. Abundance measures excluding the number of individuals for community datasets have been used in cases of large size variation among individuals within a community or where it is difficult to recognise proper individuals (Li et al., 2016). Hereafter, the Bray–Curtis indices targeting measures excluding the number of individuals (e.g. sizes and weights) are described as *d*_*BC’*_, and their components are described with apostrophes (*A’, B’*, and *C’*; detailed equations of the component see Supplementary Text 1) to distinguish them from the components in the unweighted Bray–Curtis index, as follows:

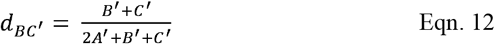

This weighting indicates that abundance-based (number of individual-based) diversity indices are weighted by total measures of individuals for each species, thus calculating diversity indices by implicitly considering the variation in the measures of each individual. As individuality is rarely clearly identifiable in weighted indices, this relationship is not well recognised.

In addition to individual-based QL indices, individual-based QT indices comprise the components of mortality and recruitment (*M* and *R*, respectively). In the QT indices, *M* refers to the total measures of individuals occurring at T*j* that died before T*k*, representing the total measures of the individuals that died between T*j* and T*k* (Fig. 1b). *R* identifies the total measures of individuals that were not yet alive at T*j* (or were not yet in the size class counted, e.g. DBH ≥ 1 cm) but were alive and counted at T*k*, representing the total measures of the individuals recruited between T*j* and T*k* (Fig. 1b).

From this point on, the major differences between the QL and QT indices are as follows. The persistent individuals are qualitatively identical, maintaining their alive (or present) status between T*j* and T*k*, but rarely maintain the same status in terms of size and weight; thus, there is a size change in an individual between T*j* and T*k*. For example, in tree communities, a newly recruited small tree can be a large tree after persisting for half a century. In the *p* component of QL indices, this implies that an individual is persistent; however, evaluating changes in its features (e.g. basal area, biomass, and/or weight) is not feasible. The indices introduced here overcome this problem by considering the contribution of changes in individual sizes, mainly targeting individual growth. Specifically, in the case of persistent individuals, the original and altered total measures are partitioned into persistent (*P*) and growth (*G*) components, respectively. Thus, the respective total measures of the components are calculated as follows:

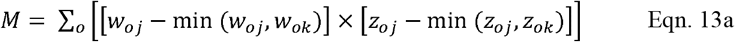

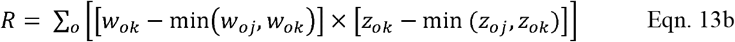

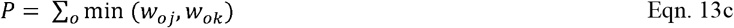

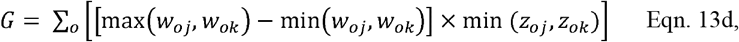

where *w*_*oj*_ is the measure of individual *o* at time T*j*, and *w*_*ok*_ is the measure of individual *o* at time T*k*. In general, basal area and biomass of each individual stem can be applicable as a measure, i.e., *w*_*oj*_ and *w*_*ok*_. *M* and *R* are the total measures of demised and recruited individuals, respectively. The latter parts of Equation 13a, b, and d identify each individual based on QL information which is equal to Equation 4a, b, and c. The persistent and growth components represent the unchanged and changed components of measures within an individual between T*j* and T*k*, respectively. Thus, *P* and *G* represent the total values of these components within a community. Equation 13d also considers decreases in the measures of each individual, because decreases in individual size may occasionally be described owing to errors, vertebrate foraging, and other unknown reasons in empirical forest datasets. Therefore, the calculation of the *G* component can be partitioned into increasing and decreasing components, as follows:

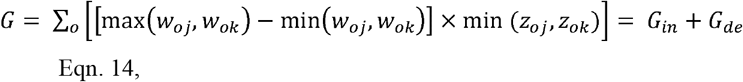

where *G*_*in*_ and *G*_*de*_ are the total increased and decreased components of the targeted measures, respectively. Specifically, the formulas for *G*_*in*_ and *G*_*de*_ are as follows:

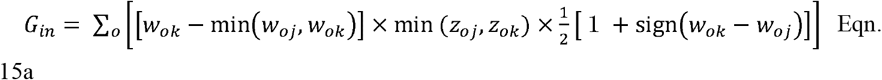

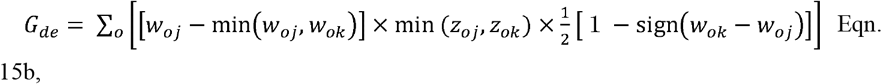

where 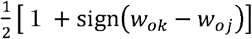 and 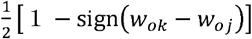 distinguish the large and small relationships (i.e. increase changes or decreased changes) between *w*_*ok*_ and *w*_*oj*_ by returning 1 only when [*w*_*ok*_ − min(*w*_*oj*_, *w*_*ok*_)] and [*w*_*oj*_ − min(*w*_*oj*_, *w*_*ok*_)] are positive values, otherwise returning 0. Additionally, min (*z*_*oj*_, *z*_*ok*_) identifies persistent individuals between time *k* and *j*, similar to that in Equation 4a.

This formulation of the individual-based QT temporal beta-diversity index for Bray– Curtis dissimilarity (*d*_*MRG*_) can be expressed as follows:

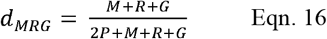

Further, this individual-based QT temporal beta diversity can be partitioned into the relative contributions of mortality (*d*_*M*_), recruitment (*d*_*R*_), and growth (*d*_*G*_) to the target weights or measurements, as follows:

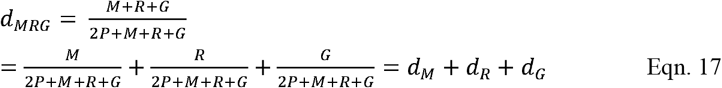

In addition, as shown in Equations 15a and 15b, the *G* component can be partitioned into *G*_*de*_ and *G*_*in*,_ which are the total decrease and increase, respectively, in the targeted measures. Equation 17 can be further partitioned into two components, as follows:

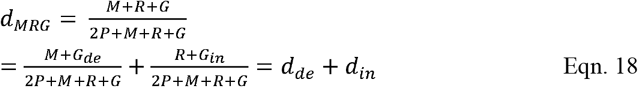

where *d*_*de*_ and *d*_*in*_ indicate the contributions of the decrease and increase, respectively, to the total changes in the targeted measures in T*j* and T*k*. In addition to the QL indices (Nakadai, 2020, 2021), the components that change over time (*M, R*, and *G*) can be reorganised into two additive components—compositional equilibrium (*E*) and shift (*B’* and *C’*). Component *M* + *G*_*de*_ can be reorganised as the total values of lost measures that contribute to compositional shifts in a community (*B’*) and the total values of lost measures replaced by conspecifics, contributing to the equilibrium in community composition (*E*_*loss*_) (Fig. 1b). Similarly, component *R + G*_*in*_ can be reorganised as the total measures that contribute to compositional shifts in a community (*C’*) and the total values of gained measures replacing conspecifics, contributing to the equilibrium in community composition (*E*_*gain*_) (Fig. 1b). By definition, the values of *E*_*loss*_ and *E*_*gain*_ are identical; hence, I replaced both *E*_*loss*_ and *E*_*gain*_ with the coefficient *E*. In addition, the persistence component (*P*) is the shared (apparent non-changed) component (*A’*) minus the community composition equilibrium (changed, but apparent non-changed) component (*E*). The above descriptions can be summarised as follows:

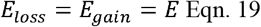

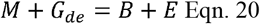

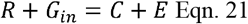

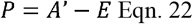

The formulation of the individual-based QT beta-diversity index for Bray–Curtis (*d*_*MRG*_) dissimilarity can be partitioned into two components—contribution to compositional shifts (*d*_*S*_) and equilibrium (*d*_*E*_)—as follows:

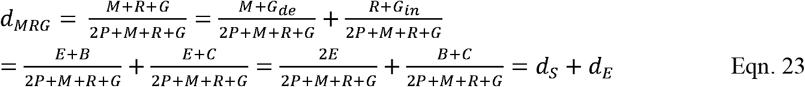

Furthermore, components *M, R*, and *G* (i.e. *M* + *G*_*de*_ and *R + G*_*in*_) can be analysed in the same manner as the community data. Therefore, the Bray–Curtis dissimilarity between communities (i.e. the total values of lost and gained measures) can be calculated based on *M, R*, and *G* (Fig. S1h), as follows:

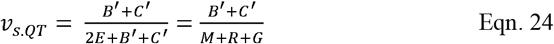

where *v*_*s.QT*_ indicates the relative speed of compositional shifts in a community against the total turnover of targeted measures associated with *M, R*, and *G*. In other words, *v*_*s.QT*_ indicates total changes in community composition divided by the total changes in targeted measures. If *v*_*s.QT*_ is 0, the turnover of the targeted measures does not contribute to compositional shifts in a community, whereas if *v*_*s.QT*_ is 1, all turnover of the targeted measures contributes to the compositional shifts. *v*_*s.QT*_ can be calculated as *d*_*BC’*_ divided by *d*_*MRG*_, as follows:

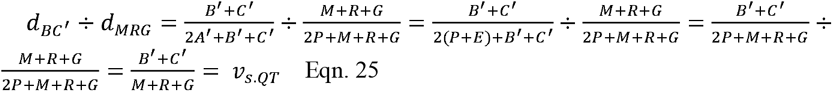

To facilitate a better understanding of the indices for potential users, I have summarised the relationships among four types of index families based on their sizes in Table S1.

Here, I focus only on an extension of Bray–Curtis dissimilarity. In Supplementary Text 2, I summarise individual-based beta diversity using the Ružička dissimilarity index (Ružička, 1958), which was identified by Podani et al. (2013) and Legendre (2014) as an appropriate index for beta-diversity studies. The Ružička dissimilarity index is an abundance-based extension of the Jaccard dissimilarity index (Jaccard, 1901).

It is noteworthy that length of the survey interval between two time periods (i.e. T*j* and T*k* in the above equations) can affect the calculated value, as well as conventional beta diversity indices (Hatosy et al., 2013), because longer intervals mean that individuals that complete a life cycle between the two periods could be lost from the investigation. In cases where a dataset with varying survey interval lengths is analysed, the length of the interval should be a sampling design-related factor.

### 2.3. Hypothetical impacts of climate warming on compositional shifts in forest communities

The hypothetical impacts of climate warming on compositional shifts, especially in forest communities, are summarised in schematic diagrams for three simple scenarios of forest compositional dynamics in Fig. 2 ((a) no climate warming, (b) compositional dynamic changes due to warming, and (c) accelerated growth and shortened life span due to warming). In scenario (b), it is assumed that the species composition of recruited individuals (i.e., changes in the species pool) become different from species in a community during the previous time step. Hence, the proportions of compositional shift (*B* and *C* in *v*_*s.QL*_ and *B’* and *C’* in *v*_*s.QT*_) increase during transitional periods. However, the total growth of individuals that have persisted since before the climate warming acts as a factor decreasing the apparent compositional shift in the size-weighted indices (*E* in *v*_*s.QT*_ increase). Scenarios (b) and (c) are represented separately to aid clarity, but they can simultaneously occur and facilitate compositional shifts in forests. In scenario (c), it is assumed that the speed of growth is twice that of scenarios (a) and (b); therefore, life spans become shorter and, thus, the turnover of both individuals and target measures increases. In contrast to scenario (b), I do not assume a change in the species pool in scenario (c), and thus, the values of both *v*_*s.QL*_ and *v*_*s.QT*_ do not change, but rather, the total turnover (*d*_*mr*_ and *d*_*MRG*_) increases. Mathematically, if *v*_*s.QL*_ and *v*_*s.QT*_ are constant, the total turnover (*d*_*mr*_ and *d*_*MRG*_) is directly linked to the apparent compositional shift (*d*_*BC*_ and *d*_*BC’*_). Individual growth largely contributes to the fluctuation in compositional shift under accelerated growth as it is more variable than individual turnover, even if life spans become shorter. Although individuals which died due to causes other than having reached the natural life span (random death and death by environmental filtering in nature) are not considered because of the complexity of visualisation in Fig. 2, in scenarios (b) and (c), such deaths may facilitate compositional shifts, specifically by increasing *v*_*s.QL*_ and *v*_*s.QT*_ or *d*_*mr*_ and *d*_*MRG*_.

**Fig. 2.**
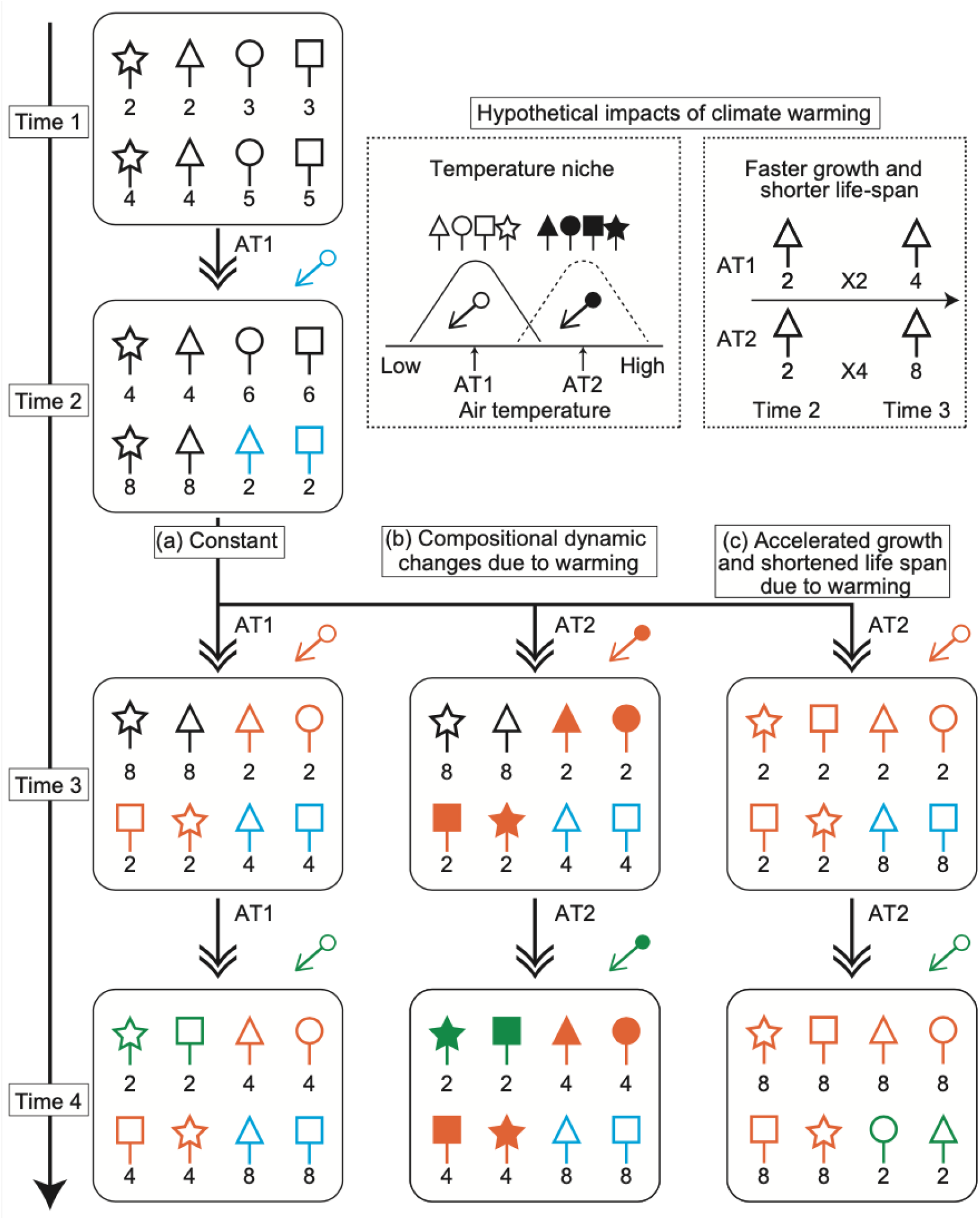
Schematic diagrams of hypothetical impacts of climate warming on compositional shifts in forest communities. Three scenarios are shown: (a) no climate warming (’Constant’, used as a comparator for the following two scenarios), (b) compositional dynamic changes due to warming, and (c) accelerated growth and shortened life span due to warming. Scenarios (b) and (c) separately show two types of potential impacts of climate warming on forest dynamics. Different symbols (both shape and fill) represent individuals of different species (i.e. eight tree species are represented here). The numbers under the symbols indicate individual size, and for simplicity, it is assumed that individuals that reach size 10 die. If an individual dies, another individual (size 2) is randomly recruited from the species pool. Two types of species pools are considered alongside air temperature differences (i.e. AT1 [no warming, constant temperature] and AT2 [warming, increase in temperature]). It is assumed that for every time step, the number doubles (e.g. from 2 to 4). The symbols of individuals in the starting communities are coloured black, while other colours indicate individuals recruited at time 2 (blue), time 3 (red), or time 4 (green). The coloured arrows with a circle (top right in each time step) indicate the type of species pool (filled or unfilled symbols). The period between time 1 and time 2 is shared among the three scenarios, and differences occur only after this period. In scenario (a) ‘constant’, it is assumed that the air temperature is constant (i.e. AT1) throughout all time periods; thus, all individuals are recruited from species indicated by unfilled symbols. Scenario (a) is included as a comparison between conventional and warming conditions. In both scenarios (b) and (c), it is assumed that the temperature increases from AT1 to AT2 after time 2. In scenario (b) ‘compositional dynamic changes’, the species pools of recruited individuals shift from unfilled-symbol species to filled-symbol species. After time 2, all recruited individuals are filled-symbol species under scenario (b); thus, until the composition completely shifts from unfilled-symbol species to filled-symbol species, the contribution of individual turnover to compositional equilibrium becomes smaller than that in the conventional condition (scenario (a)). In scenario (c) ‘accelerated growth and shortened life span’, the speed of growth increases (e.g. from 2 to 4), life spans become shorter, and the turnover of both individuals and target measures increases as a result. Note that for simplicity, deaths other than those due to having reached natural life spans are not considered.

### 2.4. Case study using subset data from the FIA database for the state of Rhode Island, USA

For a case-study application of the new individual-based QT indices based on publicly available data, I used a subset of data from the United States Department of Agriculture Forest Service’s FIA Program for the state of Rhode Island and the R package ‘rFIA’ (Stanke et al., 2020). The FIA database includes individual tree-tracking and size information (i.e. basal area and bole biomass). A comprehensive summary of the included data and sampling methodology is provided by Burrill et al. (2021). Briefly, each FIA plot comprises four 0.017-ha circular subplots, each located within a circular 0.10-ha macroplot. All free-standing woody stems (alive and dead) with a diameter at breast height (DBH) of ≥12.7 cm were sampled within each subplot (Burrill et al., 2021).

The original dataset included 10,093 individual time records for 58 species, including multiple species (spp.) in 140 plots between 2004 and 2018. For some plots, there were more than two surveys; therefore, more than one value of temporal beta diversity could be calculated. I conducted plot-level filtering of the datasets using the following criteria: (1) datasets with unidentified individuals at the species level were excluded; (2) datasets with 10 individuals or more were included; and (3) an interval of 5 ± 1 years between two surveys was targeted. In addition, individual trees with a DBH of ≥12.7 cm were targeted because bole biomass was also estimated. The final dataset included 5,705 individual time records comprising 2,541 individual trees of 32 species across 126 survey intervals (86 unique plots). The latitudinal and longitudinal ranges in the dataset were 41.39446°– 42.00797° N and 71.79635°–71.12814° W, respectively. The elevation range was 10–680 m. The geographical location of each plot is shown in Fig. S2.

I focussed on the basal area and bole biomass of each tree as changes in individual sizes that can be considered in the novel indices developed in the previous sections. The basal area of live trees was calculated as 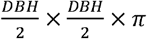. The gross cubic-foot volume of live trees was estimated at ≥12.7 cm DBH, defining the total volume of wood in the central stem of the inventoried trees (bole biomass) assuming a 30.48-centimetre mean stump height and a minimum 10.16-centimetre top diameter (for a detailed estimation method, see Domke et al., 2013).

To demonstrate the application of the novel indices, I focussed mainly on the following 11 indices: *d*_*BC*_, *d*_*mr*_, *d*_*m*_, *d*_*r*_, *v*_*s.QL*_, *d*_*BC’*_, *d*_*MRG*_, *d*_*M*_, *d*_*R*_, *d*_*G*_, and *v*_*s.QT*_. Formulas of the base indices (i.e. *d*_*BC*_, *d*_*mr*_, *v*_*s.QL*_, *d*_*BC’*_, *d*_*MRG*_, and *v*_*s.QT*_) are summarised in Table 1. All 11 indices were calculated for each survey interval for each plot. The latter six indices (i.e. *d*_*BC’*,_ *d*_*MRG*_, *d*_*M*_, *d*_*R*_, *d*_*G*_, and *v*_*s.QT*_) were calculated for both basal area and bole biomass, distinguished by the suffixes ‘_area’ or ‘_biomass’, respectively.

**Table 1.**
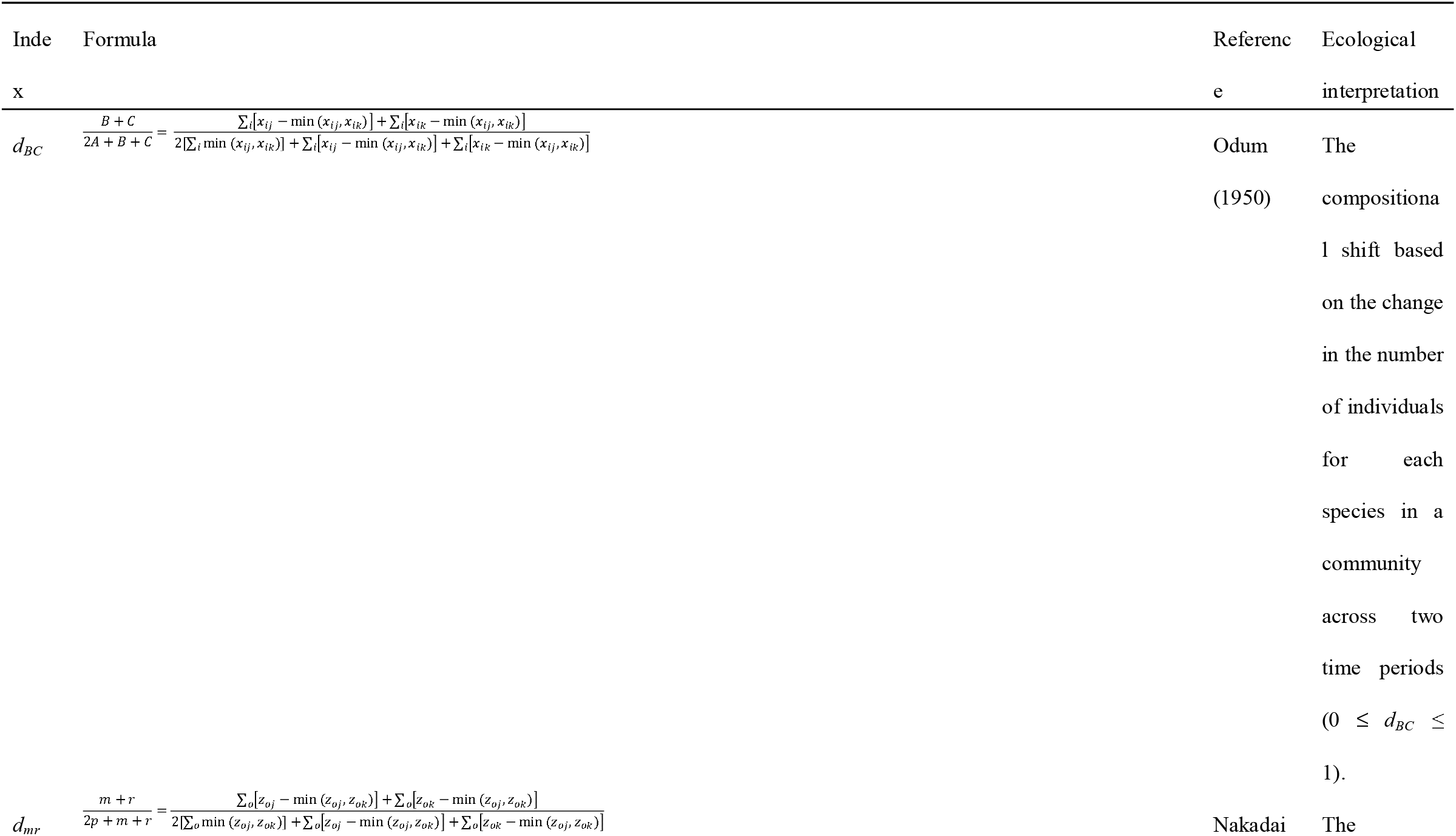
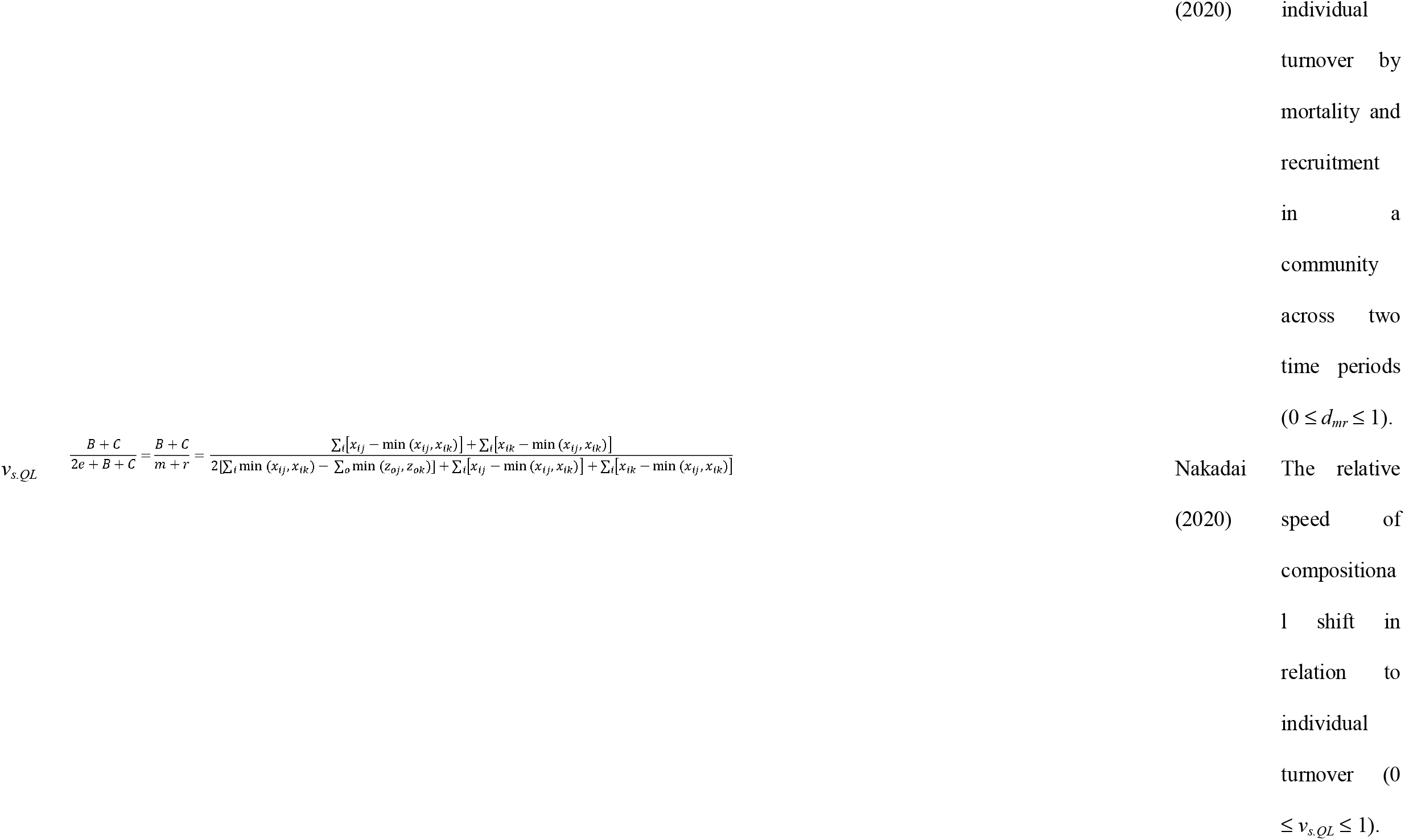
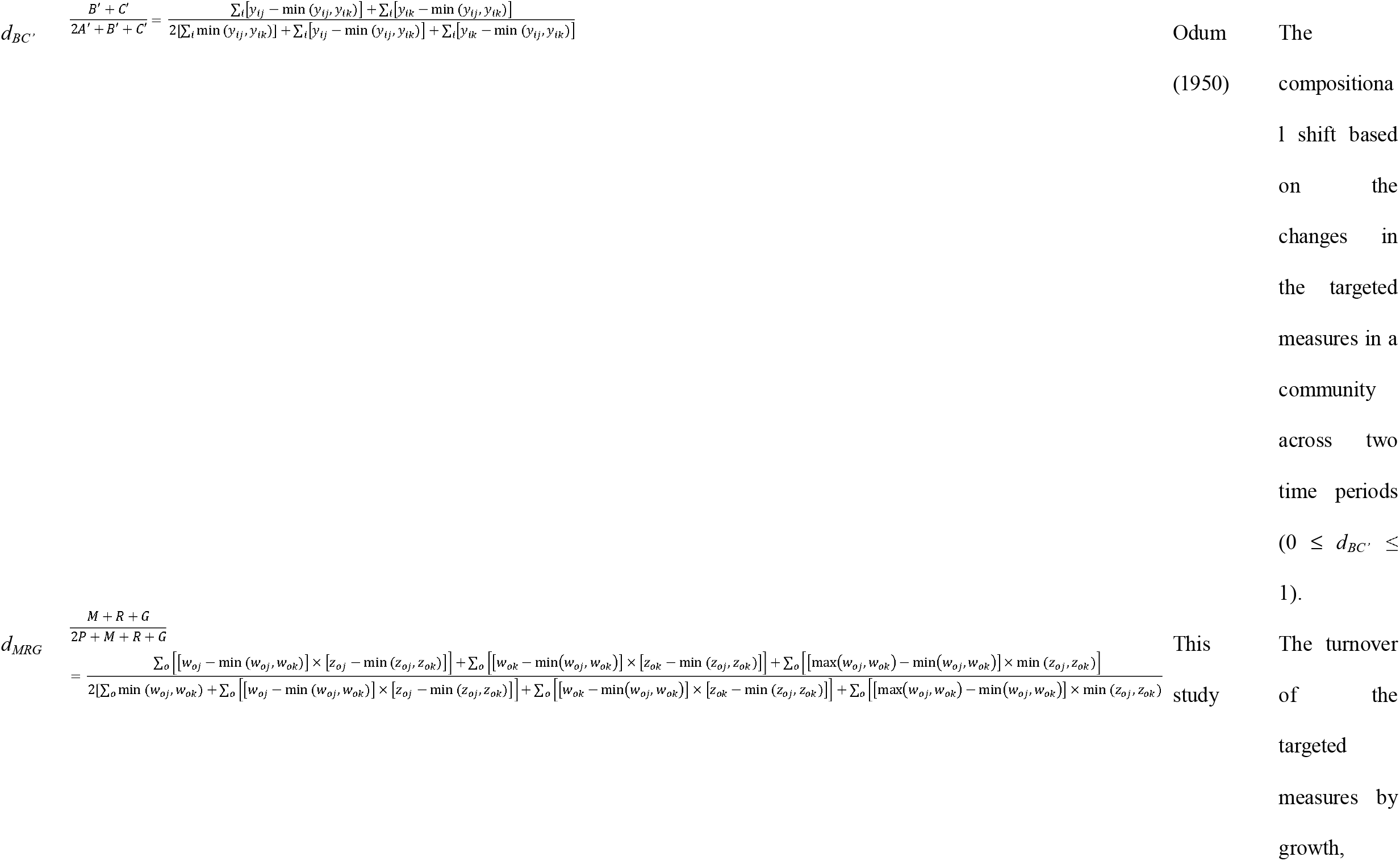
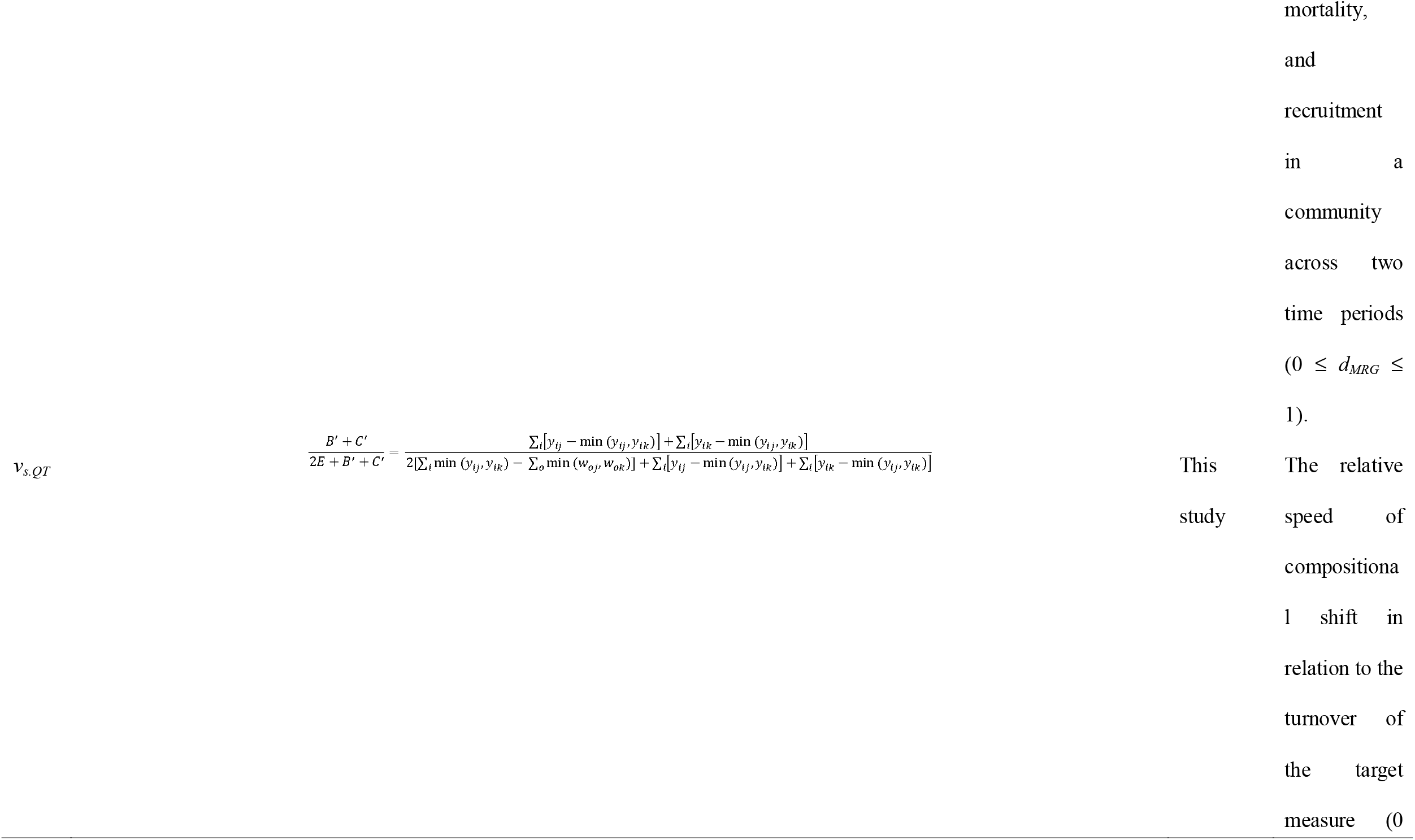
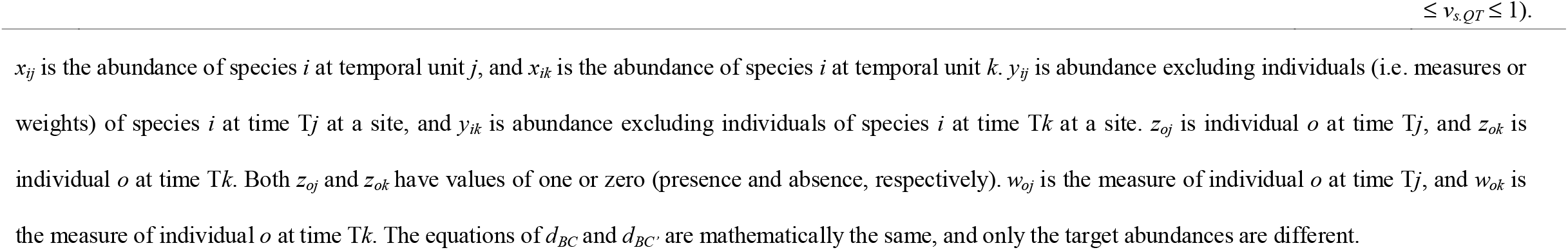
Summary of indices and formulae developed in previous studies and used in this study.

First, to show the contribution of each component to individual-based QT dissimilarity *d*_*MRG*_, I estimated kernel density for each partitioned component (*d*_*M*_, *d*_*R*_, *d*_*G*_, *d*_*S*_, and *d*_*E*_) of the dissimilarity indices. Second, I evaluated the relationships among *d*_*BC*_, *d*_*mr*,_ *v*_*s.QL*_, *d*_*BC’*_, *d*_*MRG*_, and *v*_*s.QT*_ using simple linear regression. Third, to reveal the geographical and temporal components of the indices, I ran linear mixed models for each index. As explanatory variables, geographic factors (latitude and elevation), a temporal factor (mean calendar year between the first and second surveys), and a sampling design-related factor (length of survey interval) were used. Plot identity was considered a random effect in the model comparisons because 40 plots included two survey intervals. However, model comparisons showed that the linear mixed models consistently had higher Akaike information criterion (AIC; Akaike, 1998) values than those of the linear models (Table S2). Thus, all results discussed in the following sections are based on linear models.

All of the analyses were conducted using R software (ver. 4.1.0; R Development Core Team, 2021). The R packages ‘*vegan*’ (Oksanen et al., 2020), ‘*nlme*’ (Pinheiro et al., 2022), and ‘*MuMIn*’ (Barton, 2022) were used to convert the abundance data to presence-absence data, run linear mixed models, and perform model comparisons based on AIC values. The R scripts used to calculate the new indices are provided in Supplementary File 1.

For this case study, I did not use a dataset specifically constructed for climate change assessment; hence, it is not possible to directly test the hypothetical impacts of climate warming on compositional shifts in forest communities. However, this approach helps understand the mechanisms of climate change impacts by considering changes in individual factors. Although I focus on climate change here, the indices introduced in this study can be useful in cases where individual growth rates vary over time for intrinsic reasons (e.g. ecological succession).

## 3. Results

Among the 126 survey intervals, the individual turnover of 10 survey intervals was zero; therefore, these intervals were excluded from the targets when calculating the *v*_*s*_._*QL*_ indices. The distribution of the novel indices, except *v*_*s.QT*_, is shown in Fig. 3. The contribution of individual growth to the total turnover of both basal area and bole biomass was generally larger than that of mortality and recruitment (Fig. 3a, b; Fig. S3). The *d*_*MRG*_ indices based on basal area and bole biomass were strongly and positively correlated with *d*_*BC*_, *d*_*mr*_, area-based *d*_*BC’*_, and biomass-based *d*_*BC’*_ (≥0.87; Fig. S4). In contrast, the *v*_*s.QT*_ indices based on basal area and bole biomass were only weakly correlated with the *d*_*BC’*_ indices based on basal area and bole biomass (*r* = 0.23–0.28, Fig. S4) and showed no correlation with *v*_*s.QL*_ (*r* = 0.16 and 0.08, Fig. S4). For both *d*_*MRG*_ and *v*_*s.QT*_, strong correlations existed between the indices based on basal area and bole biomass (*r* = 0.99 and 0.91, Fig. S4). All Pearson correlation coefficients are summarised in Fig. S4.

**Fig. 3.**
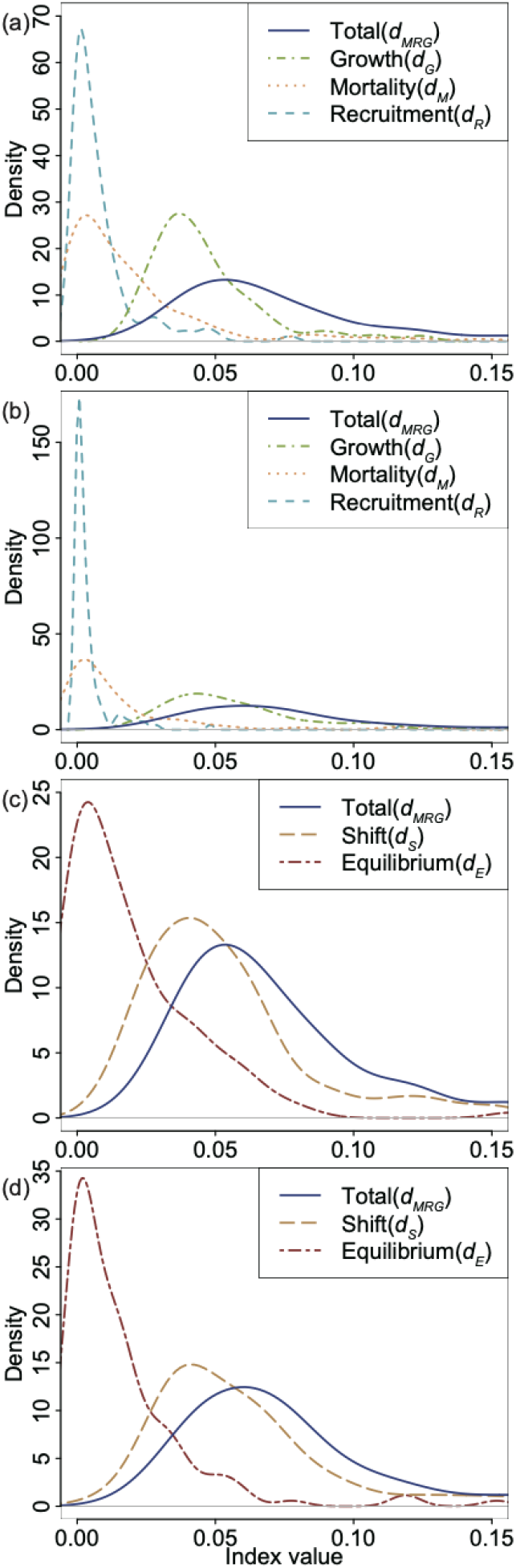
Density plots representing the distribution of individual-based QT beta-diversity indices across 126 survey intervals for 86 plots (data from Stanke et al. 2020). In all four plots, the index values are limited to the range 0–0.15 for comparison (see Fig. S3 for a plot of the entire range). The plots in panels (a) and (c) are based on basal area, and those in (b) and (d) are based on bole biomass.

Based on the AIC comparison, models including only intercept were selected for the three indices, i.e. the indices for compositional shift based on the number of individuals and basal area (*d*_*BC*_ and *d*_*BC’_area*_), and the relative speed of compositional shift against total changes in basal area (*v*_*s.QT_area*_) (Table 2). The index of compositional shift based on bole biomass (*d*_*BC’_biomass*_) tended to increase with longer survey intervals, as indicated by model comparison (Table 2). As for the index of the relative speed of compositional shift against total individual turnover (*v*_*s.QL*_), only elevation was selected as an explanatory variable based on the AIC comparison; thus, the index increased with increasing elevation (Table 2). For the relative speed of compositional shift against total changes in basal area (*v*_*s.QT_biomass*_), a model including calendar year and survey interval was selected based on the AIC comparison (Table 2); specifically, the index increased with increasing calendar year and longer survey intervals. In both indices of the total change in basal area and bole biomass (*d*_*MRG_area*_ and *d*_*MRG_biomass*_), models with latitude and survey interval were selected as the best models based on the AIC comparisons (Table 2); specifically, the indices tended to increase with decreasing latitude and longer survey intervals. However, in the index of total individual turnover (*d*_*mr*_), the model including only the survey interval was selected as the best model (Table 2), with the index increasing with longer survey intervals. As for the three indices related to the contribution of mortality to the total change (*d*_*m*_, *d*_*M_area*_, and *d*_*M_biomass*_), models including latitude and calendar year were selected as the best models (Table 2), and index values tended to increase with decreasing latitude and increasing calendar year. For the indices related to the contribution of recruitment to the total change (*d*_*R_area*_, *d*_*R_biomass*_), models including calendar year were selected as the best models (Table 2), with index values tending to increase with increasing calendar year. For the index related to the contribution of recruitment to the total individual turnover (*d*_*r*_), the model including calendar year and survey interval was selected as the best model (Table 2), with the index tending to increase with increasing calendar year and longer survey intervals. For both indices related to the contribution of growth to the total change (*d*_*G_area*_ and *d*_*G_biomass*_), the models that included the survey interval and, in the case of basal area, calendar year, were selected.

**Table 2.**
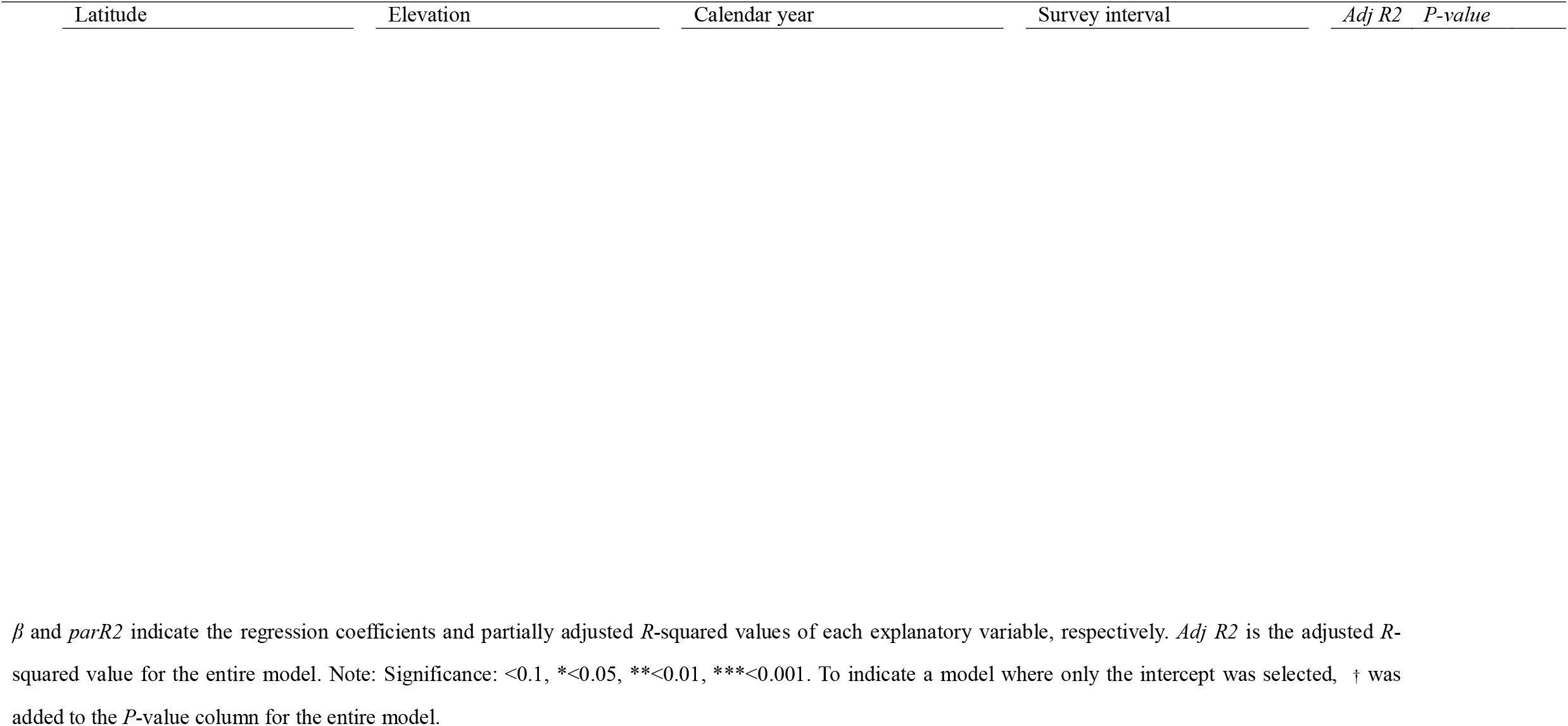
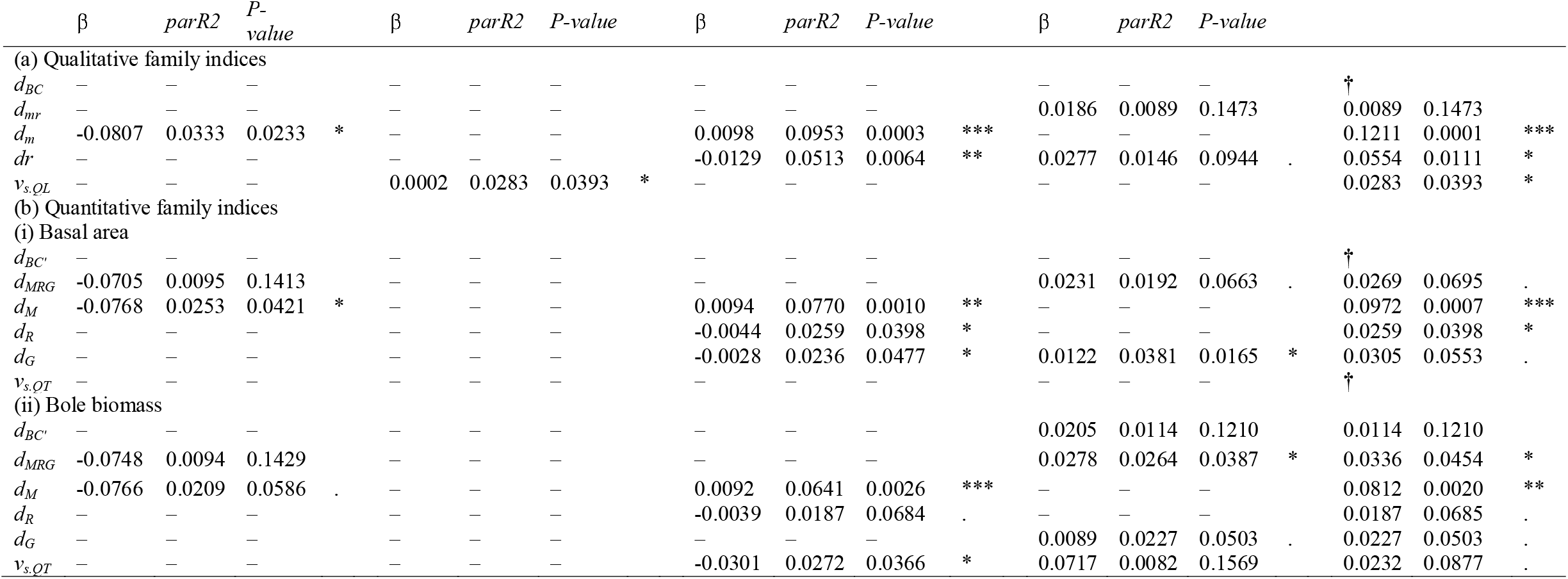
Results of a multi-regression model based on the best models selected using an Akaike information criterion (AIC) comparison for individual-based qualitative (QL) and quantitative (QT) beta-diversity indices.

## 4. Discussion

The potential for evaluating the contributions of individuals’ persistence to compositional shifts in a community was first proposed by Nakadai (2020). In this study, I extended previously established individual-based beta-diversity approaches (Nakadai, 2020, 2021) to account for individual property changes—even for persistent individuals—that are not currently considered in qualitative indices. Specifically, in addition to the components of mortality and recruitment as considered by Nakadai (2020), I introduced a growth component to account for the individual size changes in the context of compositional shifts for the first time (Fig. 1). Also, using hypothetical scenarios, I illustrated the biological meaning of the individual-based indices in the context of climate warming (Fig. 2). The novel indices proposed here not only represent methodological progress in community ecology and macroecology, but also aid conceptual advancement by bridging studies in biodiversity with those on individual life history and physiology.

To illustrate the application of the novel indices, I applied them to a subset of data from the FIA database for the state of Rhode Island, USA. The analysis demonstrated that the contribution of tree growth to the total turnover of both basal area and bole biomass was generally larger than that of mortality and recruitment (Fig. 3a, b; Fig. S3). This indicates the importance of previously overlooked growth factors in forest turnover and dynamics and the consequent compositional shifts. In the case-study dataset, the calculated abundance- and individual-based indices (i.e. *d*_*BC*_, *d*_*mr*_, *d*_*BC’*_, and *d*_*MRG*_) were strongly correlated (Fig. S4). Thus, if the contribution of individual tree growth is high, the strength of the correlation between *d*_*mr*_ and *d*_*MRG*_ should decrease, yet this was not the case. I found no clear relationships between the *v*_*s.QL*_ and *v*_*s.QT*_ indices based on both basal area and bole biomass and the relative speed of compositional shift. Generally, the *v*_*s.QL*_ values for the FIA dataset were high; thus, individual turnover generally contributed to compositional shift, although both *v*_*s.QT_area*_ and *v*_*s.QT_biomass*_ were lower and more variable than *v*_*s.QL*_. These results reflect the differences in the characteristics of the novel QT indices developed here and the conventional QL indices. Indeed, *v*_*s.QL*_ could not be applied to 10 plots that had no individual turnover, whereas in such cases, the *v*_*s.QT*_ index was more sensitive to temporal changes in the community than the *v*_*s.QL*_ index.

In the driver analysis of the calculated indices, I found that various explanatory variables affected the results (Table 2). First, focussing on the growth component specific to the new QT index, both area- and biomass-based *d*_*G*_ indices, which indicate the contribution of growth, were responsive to the survey interval. This strongly suggests that an accurate evaluation of the contribution of growth was achieved. In addition, the area-based *d*_*G*_ index tended to decrease with increasing (i.e. more recent) calendar year. In addition, overall, the contributions related to recruitment decreased, and those related to mortality increased (Table 2), which may be indicative of succession (i.e. aging forests) and/or climate change.

The *v*_*s.QT_area*_ index showed no clear pattern because the model including only intercept was selected, whereas the *v*_*s.QL*_ and *v*_*s.QT_biomass*_ indices varied with elevation and calendar year, respectively. The *v*_*s*_ family of indices is designed to capture changes to account for stochastic drifts, such as individual, basal area, or bole-volume turnover. The significant relationships between several *v*_*s*_ indices with environmental variables suggest that along with the explanatory variables, niche processes drive compositional shifts, which cannot be explained by stochastic processes alone. For example, Nakadai (2020) showed that *v*_*s.QL*_ values vary along climate gradients (i.e. the change ratio of annual mean temperature) for both deciduous and evergreen broadleaf forests.

The indices developed in the present study are applicable to classical forest inventory surveys in which all stems are tagged and their DBH measured. Here, I describe some potential applications of the novel indices as examples. First, as a representative example, applying the individual-based temporal beta-diversity indices to global datasets, especially forest inventory data (e.g. ForestGEO; https://forestgeo.si.edu/), is highly promising for assessing the impacts of global climate change. Specifically, the responses of the novel indices to both the mean and variable ratios of air temperature are important for mechanistically understanding the compositional shifts in forest tree communities. Furthermore, the average length of an individual’s life cycle (i.e. longevity) varies across the stages of ecological succession. As a base condition to study the impacts of climate change on forest tree communities, the patterns of the novel indices along the stages of ecological succession could be one of the next challenges.

Although the indices introduced in this study are mainly applicable to datasets containing both individual identity and size information, a method that accommodates various types of quantitative individual information may become available through future theoretical developments. Generally, information for individuals is limited in community ecology and macroecology, yet in the era of big data, this is likely to be addressed in the near future. In addition, if the contributions of demographic factors (i.e. growth, mortality, and recruitment) can be estimated using hierarchical modelling approaches (Hill et al., 2004, Kanamori et al., 2017), these indices could be calculated even in the absence of detailed individual-tracking information. Furthermore, although the survey interval length is treated as a sampling design-related factor in the present study, revealing the temporal patterns throughout a survey interval, and therefore the temporal scale dependency, which is called temporal distance decay (Hatosy et al., 2013), is also a future challenge.

## 5. Conclusion

Notably, climate change affects various ecological hierarchies, from global biodiversity to individuals and even intra-individual phenomena (Arneth et al., 2020; Körner, 2017). Indeed, using the newly developed individual-based temporal QT beta-diversity indices, this study showed that the growth components largely contributed to forest community compositional dynamics compared to mortality and recruitment components in the case-study dataset. The important contribution of the growth components to temporal compositional shifts (i.e. temporal beta diversity) can be evaluated using the QT indices developed in this study, thus conventional abundance-based and individual-based indices (Odam, 1950; Nakadai, 2020) cannot evaluate changes in individual sizes. Additionally, the individual-based indices can facilitate our understanding of the link between demographic processes and community compositional shifts, which conventional approaches have not achieved until now. As in the Rhode Island case study, individual-based diversity indices that account for previously overlooked elements of biodiversity, such as individual persistence and growth, pave the way for further innovative research on individual-based biodiversity. This should facilitate a greater understanding of the effects of climate change across multiple hierarchies of biological organisation.

## Author contributions

Ryosuke Nakadai: Conceptualization, Methodology, Formal analysis, Writing – original draft, review & editing.

## Funding

RN was supported by the Japan Society for the Promotion of Science (grant No. 22K15188) and the National Institute for Environmental Studies, Japan.

## Conflicts of interest

The author declares no conflicts of interest.

## Data availability

Data are available from the R package rFIA (Stanke et al., 2020), and the code used to calculate the indices in the present study is provided in Supplementary File 1.

AIC: Akaike information criterion
DBH: Diameter at breast height
FIA: Forest Inventory and Analysis

## Supplementary files

**Table S1**. Table summary to identify the applicable temporal beta-diversity indices for the community dataset

**Table S2**. Comparison of linear and linear mixed models for each index. The best models selected from the comparison are shown in Table 2

**Fig. S1**. Venn diagrams related to the calculation of both conventional and novel indices. Specifically, the first four Venn diagrams (a–d) are based on the number of individuals, and latter four (e–h) are based on the total values of targeted measures excluding the number of individuals (e.g. sizes and weights).

**Fig. S2**. Map of geographic locations of plots included in the subset of data from the Forest Inventory and Analysis (FIA) database for the state of Rhode Island, USA (Stanke et al., 2020). Sites indicated with darker-coloured points have multiple survey intervals.

**Fig. S3**. Density plots representing the distribution of individual-based quantitative beta-diversity indices across 126 survey intervals in 86 plots (data from Stanke et al., 2020). In the right four plots (b, d, f, and h), the index values are limited to the range 0–0.15 for comparison (also see Fig. 2). In the left four plots, the entire value range is shown. Plots in panels (a), (b), (e), and (f) are based on the basal area, and those in panels (c), (d), (g), and (h) on bole biomass.

**Fig. S4**. Correlations among the calculated indices. Pearson coefficients of correlation are shown. Note: Significance: <0.1, *<0.05, **<0.01, ***<0.001.

**Supplementary Text 1**. Detailed explanation for the components in the weighted Bray– Curtis index

**Supplementary Text 2**. Explanation of individual-based temporal beta diversity employing the Ružička dissimilarity index.

**Supplementary file 1**. R software script for individual-based temporal beta-diversity calculations using sample data.

## Notes

### Competing Interest Statement

The authors have declared no competing interest.

### Summary of Updates

.

## References

Akaike, H., 1998. Information theory and an extension of the maximum likelihood principle. In: E. Parzen, K. Tanabe, and G. Kitagawa (Eds.), Selected Papers of Hirotugu Akaike. Springer, New York, pp. 199–213. https://doi.org/10.1007/978-1-4612-1694-0_15.

Arneth, A., Shin, Y. J., Leadley, P., Rondinini, C., Bukvareva, E., Kolb, M., Midgley, G. F., Oberdorff T., Palomo, I., Saito, O., 2020. Post-2020 biodiversity targets need to embrace climate change. Proc. Natl. Acad. Sci. U. S. A. 117, 30882–30891. https://doi.org/10.1073/pnas.2009584117.

Bartoń, K., 2020. MuMIn: Multi-model inference. R package version 1.43.17. https://CRAN.R-project.org/package=MuMIn.

Baselga, A., 2013. Separating the two components of abundance-based dissimilarity: balanced changes in abundance vs. abundance gradients. Methods Ecol. Evol. 4, 552– 557. https://doi.org/10.1111/2041-210X.12029.

Brienen, R. J. W., Caldwell, L., Duchesne, L., Voelker, S., Barichivich, J., Baliva, M., Ceccanthini, G., Di Fillippo, A., Helama, S., Locosselli, G. M., Lopez, L. Piovesan, J., Villabla, R., Gloor, E., 2020. Forest carbon sink neutralized by pervasive growth-lifespan trade-offs. Nat. Commun. 11, 4241. https://doi.org/10.1038/s41467-020-17966-z.

Brienen, R., Phillips, O., Feldpausch, T. Gloor, E., Baker, T., Lloyd, J., Lopez-Gonzalez, G., Monteagudo-Mendoza, A., Malhi, Y., Lewis, S. L., Vásquez Martinez, R., Alexiades, M., Álvarez Dávila, E., Alvarez-Loayza, P., Andrade, A., Aragão, L. E. O. C., Araujo-Murakami, A., Arets, E. J. M. M., Arroyo, L., Aymard C. G. A., Bánki, O. S., Baraloto, C., Barroso, J., Bonal, D., Boot, R. G. A., Camargo, J. L. C., Castilho, C. V., Chama, V., Chao, K. J., Chave, J., Comiskey, J. A., Cornejo Valverde, F., da Costa, L., de Oliveira, E. A., Di Fiore, A., Erwin, T. L., Fauset, S., Forsthofer, M., Galbraith, D. R., Grahame, E. S., Groot, N., Hérault, B., Higuchi, N., Honorio Coronado, E. N., Keeling, H., Killeen, T. J., Laurance, W. F., Laurance, S., Licona, J., Magnussen, W. E., Marimon, B. S., Marimon-Junior, B. H., Mendoza, C., Neill, D. A., Nogueira, E. M., Núñez, P., Pallqui Camacho, N. C., Parada, A., Pardo-Molina, G., Peacock, J., Peña-Claros, M., Pickavance, G. C., Pitman, N. C. A., Poorter, L., Prieto, A., Quesada, C. A., Ramírez, F., Ramírez-Angulo, H., Restrepo, Z., Roopsind, A., Rudas, A., Salomão, R. P., Schwarz, M., Silva, N., Silva-Espejo, J. E., Silveira, M., Stropp, J., Talbot, J., ter Steege, H., Teran-Aguilar, J., Terborgh, J., Thomas-Caesar, R., Toledo, M., Torello-Raventos, M., Umetsu, R. K., van der Heijden, G. M. F., van der Hout, P., Guimarães Vieira, I. C., Vieira, S. A., Vilanova, E., Vos, V. A., Zagt, R. J., 2015. Long-term decline of the Amazon carbon sink. Nature 519, 344–348. https://doi.org/10.1038/nature14283.

Brown, J. H., Gillooly, J. F., Allen, A. P., Savage, V. M., West, G. B., 2004. Toward a metabolic theory of ecology. Ecology 85, 1771–1789. https://doi.org/10.1890/03-9000.

Brown, J. H., 1995. Macroecology. University of Chicago Press.

Brown, J. H., 1999. Macroecology: Progress and prospect. Oikos 87, 3–14. https://doi.org/10.2307/3546991.

Burrill, E. A., DiTommaso A. M., Turner, J. A., Pugh, S. A., Menlove, J., Christiansen, G., Perry, C. J., Conkling, B. L. 2021. The forest inventory and analysis database: Database description and user guide version 9.0.1 for Phase 2. U.S. Department of Agriculture, Forest Service, p. 1026. http://www.fia.fs.fed.us/library/database-documentation/.

Domke, G. M., Oswalt, C. M., Woodall, C. W., Turner, J. A., 2013. Estimation of merchantable bole volume and biomass above sawlog top in the National Forest Inventory of the United States. J. For. 111, 383–387. https://doi.org/10.5849/jof.13-042

Gotelli, N. J., Shimadzu, H., Dornelas, M., McGill. B., Moyes. F., & Magurran, A. E. 2017. Community-level regulation of temporal trends in biodiversity. Sci. Adv. 3, e1700315. https://doi.org/10.1126/sciadv.1700315.

Hatosy, S. M., Martiny, J. B., Sachdeva, R., Steele, J., Fuhrman, J. A., Martiny, A. C. (2013). Beta diversity of marine bacteria depends on temporal scale. Ecology 94(9), 1898–1904. https://doi.org/10.1890/12-2125.1

Hill, M. F., Witman, J. D., Caswell, H., 2004. Markov chain analysis of succession in a rocky subtidal community. Am. Nat. 164, 46–61. https://doi.org/10.1086/422340.

Jaccard, P., 1901. Distribution de la flore alpine dans le bassin des Dranses et dans quelques régions voisines. Bull. de la Soc. Vaud. des Sci. Nat. 37, 241–272.

Kanamori Y., Fukaya, K., Noda, T., 2017. Seasonal changes in community structure along a vertical gradient: Patterns and processes in rocky intertidal sessile assemblages. Popul. Ecol. 59, 301–313. https://doi.org/10.1007/s10144-017-0596-z.

Körner, C., 2017. A matter of tree longevity. Science 355, 130–131. https://doi.org/10.1126/science.aal2449.

Legendre, P., Legendre, L., 2012. Numerical ecology (3rd ed.). Amsterdam: Elsevier.

Legendre, P., 2014. Interpreting the replacement and richness difference components of beta diversity. Glob. Ecol. Biogeogr. 23, 1324–1334.

Legendre, P., 2019. A temporal beta-diversity index to identify sites that have changed in exceptional ways in space–time surveys. Ecol. Evol. 9, 3500–3514. https://doi.org/10.1002/ece3.4984.

Li, S. P., Cadotte, M. W., Meiners, S. J., Pu, Z., Fukami, T., Jiang, L., 2016. Convergence and divergence in a long-term old-field succession: The importance of spatial scale and species abundance. Ecol. Lett. 19, 1101–1109. https://doi.org/10.1111/ele.12647.

Magurran, A. E., Dornelas, M., Moyes, F., Henderson, P. A., 2019. Temporal β diversity: A macroecological perspective. Glob. Ecol. Biogeogr. 28, 1949–1960. https://doi.org/10.1111/geb.13026.

Munné-Bosch, S., 2018. Limits to tree growth and longevity. Trends Plant Sci. 23, 985– 993. https://doi.org/10.1016/j.tplants.2018.08.001.

Nakadai, R., 2020. Degrees of compositional shift in tree communities vary along a gradient of temperature change rates over one decade: Application of an individual-based temporal beta-diversity concept. Ecol. Evol. 10, 13613–13623. https://doi.org/10.1002/ece3.6579.

Nakadai, R., 2021. Individual-based multiple-unit dissimilarity: novel indices and null model for assessing temporal variability in community composition. Oecologia 197, 353–364. https://doi.org/10.1007/s00442-021-05025-3.

Odum, E. P., 1950. Bird populations of the Highlands (North Carolina) Plateau in relation to plant succession and avian invasion. Ecology 31, 587–605. https://doi.org/10.2307/1931577.

Oksanen, J., Blanchet, F. G., Friendly, M., Kindt, R., Legendre, P., McGlinn, D., Minchin, P. R., O’Hara, R. B., Simpson, G. L., Solymos, P., Stevens, M. H. H., Scöcs, E., Wagner, H. H., 2020. Vegan: Community ecology package. R package version 2.5-7. https://CRAN.R-project.org/package=vegan.

Pinheiro, J., Bates, D., DebRoy, S., Sarkar, D., R Core Team., 2022. Nlme: Linear and nonlinear mixed effects models. R package version 3.1-155. https://CRAN.R-project.org/package=nlme.

Podani, J., Ricotta, C., Schmera, D., 2013. A general framework for analyzing beta diversity, nestedness and related community-level phenomena based on abundance data. Ecol. Complex. 15, 52–61.

R Development Core Team., 2021. R: A language and environment for statistical computing. R Foundation for Statistical Computing, Vienna, Austria. (ver. 4.1.0). http://www.R-project.org/.

Ružička, M., 1958. Anwendung mathematisch–statisticher Methoden in der Geobotanik (synthetische Bearbeitung von Aufnahmen). Biologia (Bratislava) 13, 647–661.

Searle, E. B., Chen, H. Y., 2018. Temporal declines in tree longevity associated with faster lifetime growth rates in boreal forests. Environ. Res. Lett. 13, 125003. https://doi.org/10.1088/1748-9326/aaea9e.

Shimadzu, H. Dornelas, M., Magurran, A. E., 2015. Measuring temporal turnover in ecological communities. Methods Ecol. Evol. 6, 1384–1394. https://doi.org/10.1111/2041-210X.12438.

Sørensen, T. A., 1948. A method of establishing groups of equal amplitude in plant sociology based on similarity of species content and its application to analyses of the vegetation on Danish commons. Biol. Skr. 5, 1–34.

Stanke, H., Finley, A. O., Weed, A. S., Walters, B. F., Domke, G. M., 2020. rFIA: An R package for estimation of forest attributes with the US Forest Inventory and Analysis database. Environ. Model. Softw. 127, 104664. https://doi.org/10.1016/j.envsoft.2020.104664.

Tatsumi, S., Iritani, R., & Cadotte, M. W., 2021. Temporal changes in spatial variation: Partitioning the extinction and colonisation components of beta diversity. Ecol. Lett. 24(5), 1063–1072. https://doi.org/10.1111/ele.13720.

