## Supplementary figures and images for "Development of novel temporal beta-diversity indices for assessing community compositional shifts accounting for changes in the properties of individuals"

### Figure S1

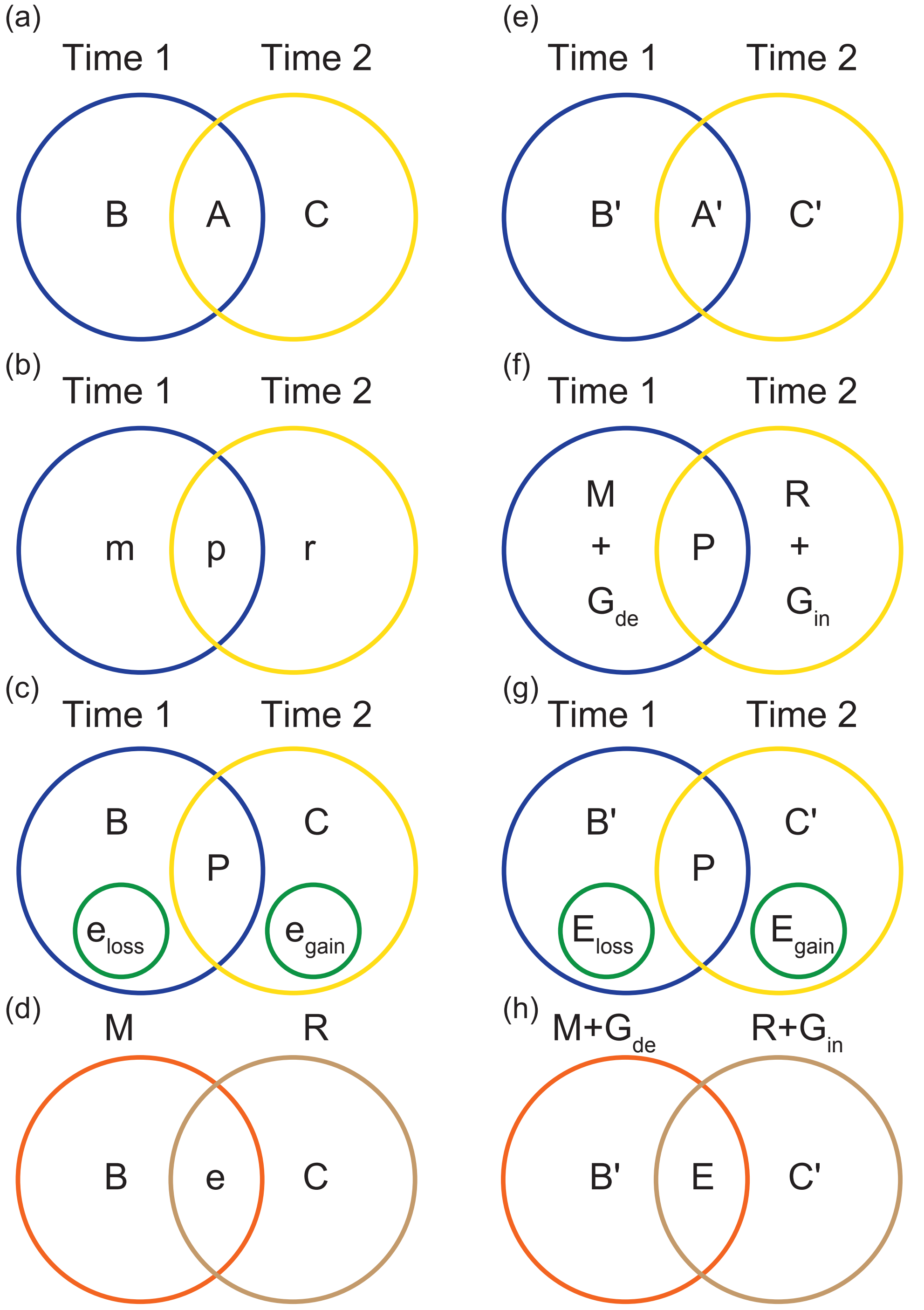

### Figure S2

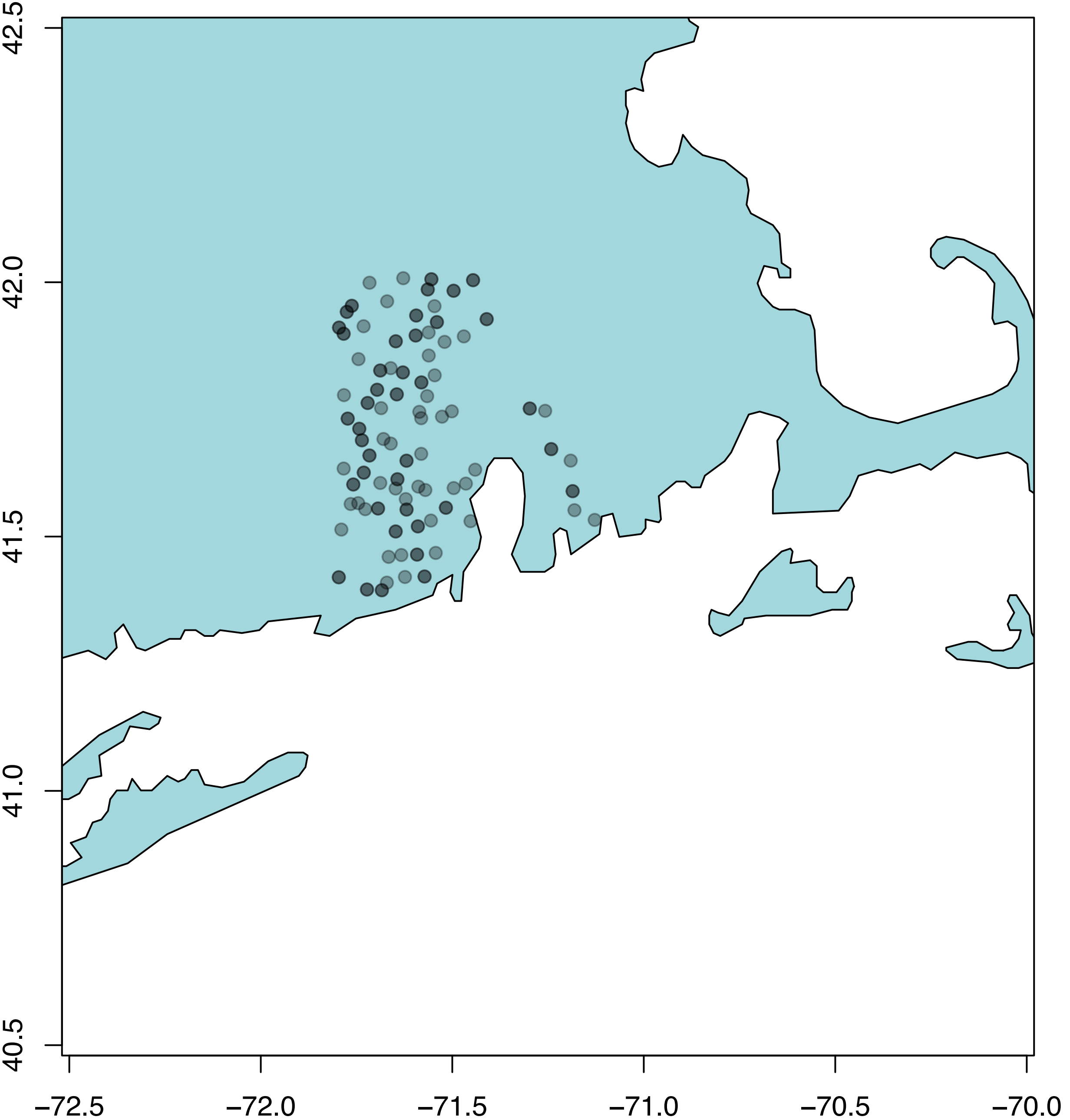

### Figure S3

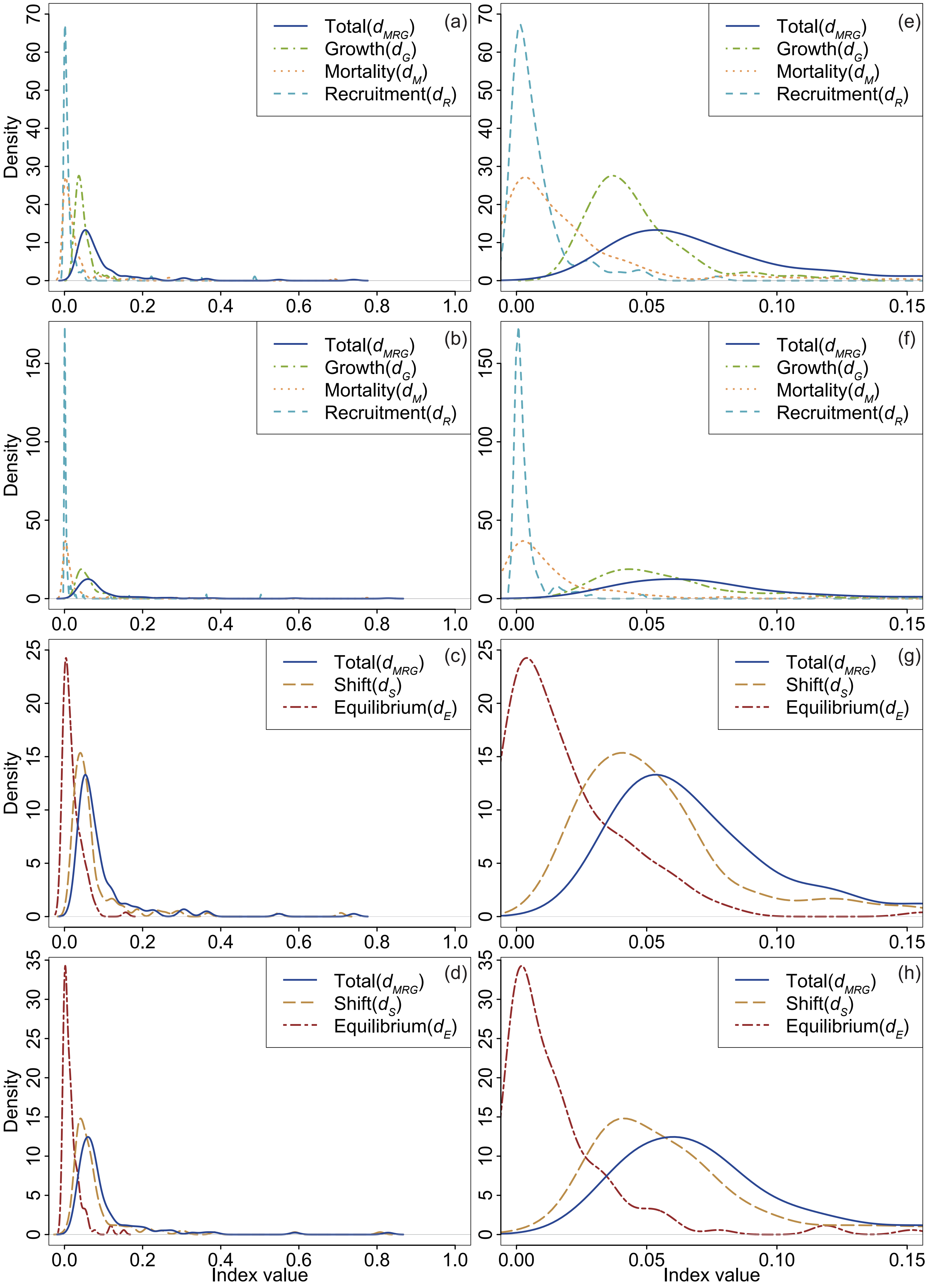

### Figure S4

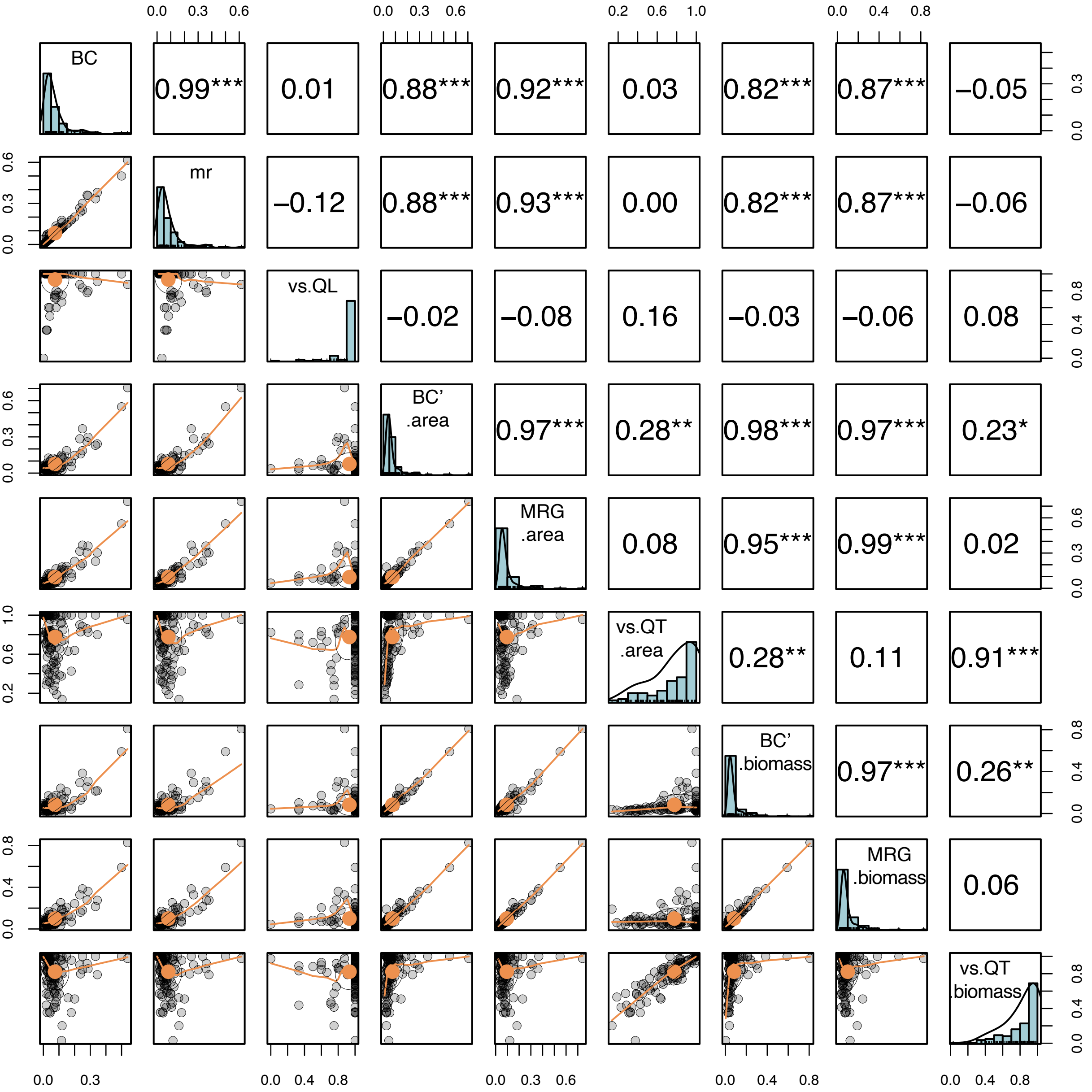
