## Supplementary material for "Development of novel temporal beta-diversity indices for assessing community compositional shifts accounting for changes in the properties of individuals": Table S1

Table S1. Table summary to identify the applicable temporal beta-diversity indices for the community dataset.

|  |  | Is community data based on countable number of individuals or total measures (e.g. total basal area) for each species as unit? | | |
| --- | --- | --- | --- | --- |
|  |  | countable number of individuals | only total measures | Both |
| Does target community data include individual identity information? | Yes | 1, 3 | 2, 4 | 1, 2, 3, 4 |
|  | No | 1 | 2 | 1, 2 |

Note: The numbers correspond to the indices (1. Bray‒Curtis dissimilarity index based on the number of individuals, 2. Bray‒Curtis dissimilarity index based on a specific measure, 3. individual-based QL beta-diversity indices, and 4. individual-based QT beta-diversity indices).
